# The metabolic response of *Pseudomonas taiwanensis* to NADH dehydrogenase deficiency

**DOI:** 10.1101/624536

**Authors:** Salome C. Nies, Robert Dinger, Yan Chen, Gossa G. Wordofa, Mette Kristensen, Konstantin Schneider, Jochen Büchs, Christopher J. Petzold, Jay D. Keasling, Lars M. Blank, Birgitta E. Ebert

**Affiliations:** iAMB-Institute of Applied Microbiology, ABBt-Aachen Biology and Biotechnology, RWTH Aachen University, DE; AVT – Biochemical Engineering, RWTH Aachen University, DE; Joint BioEnergy Institute, Emeryville, CA 94608, USA; Novo Nordisk Foundation Center for Biosustainability, Technical University of Denmark, DK-2800 Lyngby, DK; Lawrence Berkeley National Laboratory, Biological Systems and Engineering Division, Berkeley, CA 94720, USA; Virtual Institute of Microbial Stress and Survival, Lawrence Berkeley National Laboratory, Berkeley, CA; Physical Biosciences Division, Lawrence Berkeley National Laboratory, Berkeley, CA; Dept. of Bioengineering, University of California, Berkeley, CA; Dept. of Chemical Engineering, University of California, Berkeley, CA; Synthetic Biochemistry Center, Institute for Synthetic Biology, Shenzhen Institutes for Advanced Technologies, Shenzhen, China

**Keywords:** *Pseudomonas*, NADH dehydrogenase, respiratory activity, oxidative stress, electron transport chain

## Abstract

Obligate aerobic organisms rely on a functional electron transport chain for energy generation and NADH oxidation. Because of this essential requirement, the genes of this pathway are likely constitutively and highly expressed to avoid a cofactor imbalance and energy shortage under fluctuating environmental conditions.

We here investigated the essentiality of the three NADH dehydrogenases of the respiratory chain of the obligate aerobe *Pseudomonas taiwanensis* VLB120 and the impact of the knockouts of corresponding genes on its physiology and metabolism. While a mutant lacking all three NADH dehydrogenases seemed to be nonviable, the generated single or double knockout strains displayed none or only a marginal phenotype. Only the mutant deficient in both type 2 dehydrogenases showed a clear phenotype with biphasic growth behavior and strongly reduced growth rate in the second phase. In-depth analyses of the metabolism of the generated mutants including quantitative physiological experiments, transcript analysis, proteomics and enzyme activity assays revealed distinct responses to type II and type I dehydrogenase deletions. An overall high metabolic flexibility enables *P. taiwanensis* to cope with the introduced genetic perturbations and maintain stable phenotypes by rerouting of metabolic fluxes.

This metabolic adaptability has implications for biotechnological applications. While the phenotypic robustness is favorable in large-scale applications with inhomogeneous conditions, versatile redirecting of carbon fluxes upon genetic interventions can frustrate metabolic engineering efforts.

**Importance:** While *Pseudomonas* has the capability for high metabolic activity and the provision of reduced redox cofactors important for biocatalytic applications, exploitation of this characteristic might be hindered by high, constitutive activity of and consequently competition with the NADH dehydrogenases of the respiratory chain. The in-depth analysis of NADH dehydrogenase mutants of *Pseudomonas taiwanensis* VLB120 presented here, provides insight into the phenotypic and metabolic response of this strain to these redox metabolism perturbations. The observed great metabolic flexibility needs to be taken into account for rational engineering of this promising biotechnological workhorse towards a host with controlled and efficient supply of redox cofactors for product synthesis.

## Introduction

Many industrially relevant molecules, e. g., ethanol, butanediol or isoprene, are more reduced than the industrially-used sugars glucose and sucrose or alternative, upcoming carbon sources such as xylose or glycerol (1–3). The microbial production of those favored compounds hence is inherently redox limited, i.e. by the supply of reduced redox cofactors, generally NADH or NADPH. This bottleneck has been overcome in some cases, e.g., 1,4 butanediol and 1,3-propanediol production in *Escherichia coli* (4, 5) or L-lysine synthesis in *Corynebacterium glutamicum* (6). Applied strategies are optimization of the host metabolism by metabolic engineering (4, 7, 8) or adaptation of process conditions by (co-)feeding reduced substrates (9), applying microaerobic conditions or by nongrowing cells with reduced competition and cellular demand for the redox cofactor (10–13). Alternatively, microorganisms can be applied that naturally outperform the classic, industrial workhorses with respect to redox cofactor supply. Pseudomonads are outstanding in this regard as they exhibit a driven-by-demand phenotype, which allows them to strongly enforce the metabolic activity under stress conditions with increased energy demand, reported to result in a more than 2-fold carbon uptake rate and an even 8-fold increase of the NAD(P)H regeneration rate relative to standard growth conditions (12, 14, 15). This behavior holds great promise for using this species for the bioproductions of highly reduced chemicals such as phenol, (*S*)-styrene oxide, rhamnolipids, and methyl ketones (16–20). Yet, competition is high as the NAD^+^/NADH couple functions as cofactor in over 300 oxidation/reduction reactions (21). The obligate aerobic lifestyle of *Pseudomonas* without apparent fermentative metabolism necessitates constitutive activity of the NADH dehydrogenases to ensure adequate oxidation of NADH to NAD^+^. Hence, we argue here that a naturally high NADH oxidation activity might impair the effective fueling of production pathways with reducing equivalents. We here set out to provide an in-depth analysis of the redox metabolism of *Pseudomonas taiwanensis* VLB120, a strictly aerobic bacterium, focusing on the role and essentiality of the single NADH dehydrogenases for NADH oxidation and energy generation.

The genome of *P. taiwanensis* VLB120 encodes two types of NADH dehydrogenases, type I (EC 7.1.1.2) and two isoforms of type II (EC 1.6.99.3). Type I is encoded by the genes PVLB_15600-15660 designated as the *nuo* operon and often referred to as complex 1 (22). Nuo is a multisubunit enzyme complex and couples the electron transfer to proton translocation (23). The resulting proton gradient can then be used by the ATP synthase for the generation of ATP. The two type II NADH dehydrogenases, also termed alternative NADH dehydrogenase, are encoded by PVLB_13270 and PVLB_21880, designated as *ndh1* and *ndh2*, respectively. Ndh1 and Ndh2, both consist of a single polypeptide chain; they transfer electrons from NADH to ubiquinone but do not contribute to the membrane potential (23, 24). The amino acid sequence of the NADH dehydrogenases *ndh2* and *nuoA-N* of *P. taiwanensis* VLB120 and *Pseudomonas putida* KT2440 share a 96% homology.

In this study, NADH dehydrogenase mutants of *P. taiwanensis* VLB120 were generated and characterized regarding growth, respiratory activity, and transcriptional and proteomic changes to elucidate the impact of redox metabolism perturbation on the cellular physiology.

## Materials and Methods

### Strains, media and culture conditions

Bacterial strains used in this study are listed in Table 1. Strains were propagated in Lysogeny Broth (LB) containing 10 g L^−1^ peptone, 5 g L^−1^ sodium chloride, and 5 g L^−1^ yeast extract (25). Cetrimide agar (Sigma-Aldrich, St. Louis, MO, USA) was used after mating procedures to select for *Pseudomonas*. Growth and characterization experiments were performed using mineral salt medium (MSM) (26) containing 3.88 g L^−1^ K_2_HPO_4_, 1.63 g L^−1^ NaH_2_PO_4_, 2 g L^−1^ (NH_4_)_2_SO_4_, 0.1 g L^−1^ MgCl_2_·6H_2_O, 10 mg L^−1^ EDTA, 2 mg L^−1^ ZnSO_4_·7 H_2_O, 1 mg L^−1^ CaCl_2_·2H_2_O, 5 mg L^−1^ FeSO_4_·7 H_2_O, 0.2 mg L^−1^ Na_2_MoO_4_·2H_2_O, 0.2 mg L^−1^ CuSO_4_·5 H_2_O, 0.4 mg L^−1^ CoCl_2_·6 H_2_O, 1 mg L^−1^ MnCl_2_·2 H_2_O supplemented with 25 mM glucose. For preparation of solid LB, 1.5% agar was added to the medium. For plasmid maintenance and in the gene deletion procedure, antibiotics were added to the medium as required. Gentamycin and Kanamycin were used at concentrations of 25 mg L^−1^ and 50 mg L^−1^, respectively.

**Table 1.** Bacterial strains used in this study.

| Strain | Relevant characteristics<sup>a</sup> | Reference |
| --- | --- | --- |
| ***E. coli*** |  |  |
| DH5α | *fhuA2 lac(del)U169 phoA glnV44*<br>Φ80' *lacZ(del)M15 gyrA96 recA1*<br>*relA1 endA1 thi-1 hsdR17* | NEB |
| DH5αλpir1 | F<sup>-</sup>, Δ*lac*169, *rpoS*(Am), *robA1*,<br>*creC510*, *hsdR514*, *endA*,<br>*recA1uidA(ΔMluI)::pir-116*; host for<br>*oriV*(R6K) vectors in high copy<br>number | Thermo Fisher<br>Scientific |
| HB101 pRK2013 | Sm<sup>R</sup>, *hsdR-M+*, *proA2*, *leuB6*, *thi-1*,<br>*recA*; bears plasmid pRK2013 | (27) |
| DH5α pSW-2 | Gm<sup>r</sup>, DH5α bearing pSW-2 | (28) |
| DH5αλpir1 pEMG | Km<sup>r</sup>, DH5αλpir1 bearing plasmid<br>pEMG | (28) |
| DH5αλpir1<br>pEMG_ko_ndh1 | Km<sup>r</sup>, PVLB_13270 deletion plasmid | This study |
| DH5αλpir1<br>pEMG_ko_ndh2 | Km<sup>r</sup>, PVLB_21880 deletion plasmid | This study |
| DH5αλpir1<br>pEMG_ko_nuo | Km<sup>r</sup>, PVLB_15600-15660 deletion<br>plasmid | This study |
| ***P. taiwanensis*** |  |  |
| VLB120 | wild type | (29) |
| VLB120 pSTY<sup>-</sup> | VLB120 devoid of megaplasmid<br>pSTY | Wynands et al.<br>submitted |
| VLB120 Δ*ndh1* | Δ*ndh1* (PVLB_13270), pSTY<sup>-</sup> | This study |
| VLB120 Δ*ndh2* | Δ*ndh2* (PVLB_21880), pSTY<sup>-</sup> | This study |
| VLB120 ΔΔ*ndh* | ΔΔ*ndh* (PVLB_13270, PVLB_21880),<br>pSTY<sup>-</sup> | This study |
| VLB120 Δ*nuo* | Δ*nuo* (PVLB_15600-15660), pSTY<sup>-</sup> | This study |
| VLB120 Δ*nuo*Δ*ndh1* | Δ*ndh1* (PVLB_13270), Δ*nuo*<br>(PVLB_15600-15660), pSTY<sup>-</sup> | This study |

<sup>a</sup> Gm<sup>r</sup>, Km<sup>r</sup>, gentamycin, kanamycin resistance, respectively.

Batch flask experiments were performed in 50 mL medium in 500-mL flasks on a horizontal rotary shaker with a throw of 50 mm and frequency of 300 rpm. *E. coli* was grown at 37 °C, *Pseudomonas* at 30 °C. The chemicals used in this work were obtained from Carl Roth (Karlsruhe, Germany), Sigma-Aldrich (St. Louis, MO, USA), or Merck (Darmstadt, Germany), unless stated otherwise. The main cultures were inoculated from liquid pre-cultures to an approximate OD_600nm_ of 0.05. All experiments were performed in biological triplicates unless stated otherwise.

### Plasmid cloning and generation of deletion strains

Genomic DNA of *P. taiwanensis* VLB120 was isolated using the High Pure PCR Template Preparation Kit (Hoffmann-La-Roche, Basel, Switzerland). Upstream (TS1) and downstream (TS2) regions with a length of 400-800 bp flanking the specific target gene were amplified using Q5 High-Fidelity Polymerase (New England Biolabs, Ipswich, MA, USA). Primers were ordered as unmodified DNA Oligonucleotides from Eurofins Genomics (Ebersberg, Germany) and are listed in Supplementary Table S1. The suicide delivery vector pEMG was isolated using the NEB Monarch Plasmid Miniprep Kit (New England Biolabs, Ipswich, MA, USA). The isolated plasmid was digested with restriction enzymes purchased from New England Biolabs (Ipswich, MA, USA). For plasmid construction, Gibson Assembly using NEB Builder Hifi DNA Assembly (New England Biolabs, Ipswich, MA, USA) was used. Plasmids were transformed into electrocompetent *E. coli* DH5αλpir1 via electroporation (30). Transformants and chromosomally engineered *Pseudomonas* were screened by colony PCR using OneTaq 2x Master Mix (New England Biolabs, Ipswich, MA, USA). The cell material was lysed in alkaline polyethylene glycol for enhanced colony PCR efficiency as described previously (31).

Targeted gene deletions were performed using the I-SceI-based system developed by Martinez-Garcia and de Lorenzo (28). The conjugational transfer of the mobilizable knock-out plasmids from *E. coli* DH5αλpir1 to *Pseudomonas* was performed via triparental patch mating (16). After conjugation, the pSW-2 plasmid encoding the I-SceI-endonuclease was conjugated into *Pseudomonas* co-integrates. The addition of 3-methylbenzoate for the induction of I-SceI expression was omitted as the basal expression level was sufficient. Kanamycin-selective clones were directly isolated, positive clones were cured of pSW-2 and restreaked several times. The gene deletion was confirmed by colony PCR and Sanger sequencing.

### Analytical methods

The optical density of cell suspensions was measured at a wavelength of 600 nm using an Ultrospec 10 spectrophotometer (GE Healthcare, Chicago, IL, USA). The cell dry weight (CDW) was calculated by multiplying OD_600nm_ with a gravimetrically determined correlation factor of 0.39. For HPLC analysis the samples were centrifuged at 17,000 x g for 5 min and the supernatant was stored at −20°C until further analysis.

Glucose and gluconate concentrations were measured by high-performance liquid chromatography using a Beckman System Gold 126 Solvent Module equipped with a System Gold 166 UV-detector (Beckman Coulter) and a Smartline RI detector 2300 (KNAUER Wissenschaftliche Geräte, Berlin, Germany). Analytes were separated on the organic resin column Metab AAC (ISERA, Düren, Germany) eluted with 5 mM H_2_SO_4_ at an isocratic flow of 0.6 mL min^−1^ at 40 °C for 20 min. Glucose and gluconate were analyzed using the RI detector whereas gluconate was determined with the UV detector at a wavelength of 210 nm.

### Proteomic profiling of NADH dehydrogenase mutants

Samples for proteome profiling were taken during early-, mid-, and late-exponential growth at an OD_600nm_ of 0.5, 2.5, and after depletion of glucose, checked with test strips for rapid detection of glucose (Medi-Test, Macherey-Nagel, Düren, Germany), respectively. Proteins were extracted from cell biomass and subsequently prepared for shotgun proteomic experiments as described previously (32). All samples were analyzed on an Agilent 6550 iFunnel Q-TOF mass spectrometer (Agilent Technologies, Santa Clara, CA) coupled to an Agilent 1290 UHPLC system. Twenty (20) μg of peptides were separated on a Sigma–Aldrich Ascentis Peptides ES-C18 column (2.1 mm × 100 mm, 2.7 μm particle size, operated at 60°C) at a 0.400 mL min^−1^ flow rate and eluted with the following gradient: initial condition was 95% solvent A (0.1% formic acid) and 5% solvent B (99.9% acetonitrile, 0.1% formic acid). Solvent B was increased to 35% over 120 min, and then increased to 50% over 5 min, then up to 90% over 1 min, and held for 7 min at a flow rate of 0.6 mL min^−1^, followed by a ramp back down to 5% B over 1 min where it was held for 6 min to re-equilibrate the column to original conditions. Peptides were introduced to the mass spectrometer from the LC by using a Jet Stream source (Agilent Technologies) operating in positive-ion mode (3,500 V). Source parameters employed gas temp (250°C), drying gas (14 L min^−1^), nebulizer (35 psig), sheath gas temp (250°C), sheath gas flow (11 L min^−1^), VCap (3,500 V), fragmentor (180 V), OCT 1 RF Vpp (750 V). The data were acquired with Agilent MassHunter Workstation Software, LC/MS Data Acquisition B.06.01 operating in Auto MS/MS mode whereby the 20 most intense ions (charge states, 2–5) within 300–1,400 m/z mass range above a threshold of 1,500 counts were selected for MS/MS analysis. MS/MS spectra (100–1,700 m/z) were collected with the quadrupole set to “medium” resolution and were acquired until 45,000 total counts were collected or for a maximum accumulation time of 333 ms. Former parent ions were excluded for 0.1 min following MS/MS acquisition. The acquired data were exported as mgf files and searched against the pan proteome that is highly related to *Pseudomonas taiwanensis* VLB120 with Mascot search engine version 2.3.02 (Matrix Science). The resulting search results were filtered and analyzed by Scaffold v 4.3.0 (Proteome Software Inc.). The normalized spectra count of each sample were exported from Scaffold, and the relative quantity changes of identified proteins in mutant samples were calculated in comparison to wild type sample. The statistical significance of these changes and the adjusted p-values were evaluated by limma R package. The mass spectrometry proteomics data have been deposited to the ProteomeXchange Consortium via the PRIDE (33) partner repository with the dataset identifier PXD013623 and 10.6019/PXD013623.

### RNA preparation and analysis

Samples for transcription analysis were taken during early-, mid- and late-exponential growth at an OD_600nm_ of approximately 0.5, approximately 2.5 and after glucose depletion, respectively. Prior to RNA isolation, the culture sample was diluted with the DNA/RNA protection reagent of the Monarch Total RNA Miniprep Kit (New England Biolabs, Ipswich, MA, USA), followed mechanical lysis with ZR BashingBead™ Lysis Tube (0.5mm) (Zymo Research, Irvine, CA, USA) for 1 min using the Mini-Beadbeater-16 (Biospec, Bartlesville, OK, USA). After a centrifugation step at 16,000 x g for 2 min the supernatant was transferred into a new tube. An equal volume of RNA lysis buffer of the Monarch Total RNA Miniprep Kit was added, and the RNA isolation was continued as described in the supplier’s manual. After the last elution step, an additional in-tube DNase treatment was done using RNase-free DNaseI (New England Biolabs, Ipswich, MA, USA). The final RNA yield and purity were evaluated by the absorption ratio A_260_/A_280_ measured with a Nanodrop (Thermo Scientific, Rockford, IL, USA). The synthesis of cDNA for reverse transcription was carried out with a Protoscript II first strand cDNA synthesis kit (New England Biolabs, Ipswich, MA, USA) using 120 ng total RNA and 60 μM random hexamers. The qPCR analyses were conducted with 5 μL of the reverse transcription reaction mixture with gene-specific primers (Supplementary Table S1) and the Luna Universal qPCR Master Mix (New England Biolabs, Ipswich, MA, USA) was used. Primers for qPCR were designed with the PrimerQuest Tool of IDT technologies. Gene expression levels for each individual sample were normalized relative to the internal reference gene, *rpoB* and the wild type in the corresponding growth phase calculated by a mathematical method based on the calculated real-time PCR efficiencies (34). The qPCR was performed with the CFX96 Real-Time PCR Detection System (Biorad, Hercules, CA, USA). All qPCR reactions were performed in triplicates.

### Inverted membrane vesicle preparation and NADH oxidation activity

Cultures were harvested at early-exponential growth phase at an optical density (OD_600nm_) of approximately 0.5, as well as in the late-exponential growth phase (OD_600nm,_ of 3-4). Inverted membrane vesicles were prepared as described by Borisov (35). Briefly, cells were centrifuged for 8 min at 5,000 x g and resuspended in 2 mL spheroblast buffer (200 mM Tris-HCl pH 8.0, 2 mM EDTA, 30% sucrose), centrifuged again and resuspended in 1 mL spheroplast buffer. Spheroplasts were prepared using lysozym (0.03 g) and incubated for 30 min at room temperature. Spheroplasts were centrifuged for 10 min at 5,000 x g and resuspended in 2 mL sonication buffer (100 mM HEPES-KOH pH 7.5, 50 mM K_2_SO_4_, 10 mM MgSO_4_, 2 mM DTT, 0.5 mM PMSF). The vesicles were sonicated (Bioruptor, Diadenode, Belgium) in 4 cycles à 30 sec at high intensity with an intermediate pause of 30 sec in ice water. The inverted membrane vesicles were centrifuged twice for 10 min at 5,000 x g to remove cell debris. The supernatant was centrifuged for 30 min at 120,000 x g and the resulting pellet was resuspended in the assay buffer (25 mM HEPES, 25 mM BIS-TRIS propane pH 7, 10 mM MgSO_4_).

The freshly prepared inverted membrane vesicles were immediately used for the determination of the NADH oxidation activity as we observed a rapid activity decline when the membrane samples were stored on ice. 150 μL mL^−1^ isolated membrane fractions were added to the assay buffer and the reaction was initiated by the addition of 125 μM NADH, the total volume of the assay was 200 μL. The NADH oxidation was monitored over 30 min at 340 nm in a Synergy™ MX microplate reader (BioTek, Winooski, VT, USA). For calculating the specific enzyme activity, we used the NADH molar extinction coefficient ԑ_NADH_ = 6.22 mM^−1^ cm^−1^; one unit of activity was the quantity that catalyzed the oxidation of 1 μmol of NADH per min. The protein concentration was measured with the reducing agent compatible Pierce BCA Protein Assay Kit (Thermo Scientific, Rockford, IL, USA).

### Respiration activity monitoring

The cultivations and measurements of the oxygen transfer rate (OTR) and the carbon dioxide transfer rate (CTR) were performed in a modified RAMOS System, developed by the Chair of Biochemical Engineering (RWTH Aachen University) (36, 37). The standard RAMOS for shake flasks is commercially available from the Kühner AG (Birsfelden, Switzerland) or HiTec Zang GmbH (Herzogenrath, Germany). All cultivations were performed in 250-mL Ramos flasks with 10 % (v/v) filling volume using MSM medium supplemented with 25 mM glucose. The cultures were inoculated from liquid pre-cultures to an approximate OD_600nm_ of 0.05. The OTR and CTR were measured thrice per hour. All experiments were performed in biological duplicates.

### Redox cofactor quantification

Samples for redox cofactor analysis were collected from early-, mid-, and late-exponential growth phase at OD_600nm_ of approximately 0.6, 2.2 and 4.0, respectively. The samples were rapidly transferred into a 15-mL falcons tube containing 5 mL of quenching solution (acetonitrile:methanol:water, 40:40:20, v/v) with ^13^C labelled cell extracts at –40°C.

After three freeze-thaw cycles, the samples were centrifuged at 13,000 x g for 5 min and concentrated by evaporating the quenching solvent using a vacuum concentrator (SAVANT, SpeedVac, Thermo Fisher Scientific, San Diego, CA, USA) for 5 hours followed by lyophilization (LABCONCO, FreeZone, Kansas City, MO, USA). All dried extracts were stored −80°C until analysis or re-suspended in LC-MS grade water for LC-MS analysis.

All redox cofactor metabolites were measured on an AB SCIEX Qtrap1 5500 mass spectrometer (AB SCIEX, Framingham, MA, USA) operated in negative ion and selected multiple reaction monitoring (MRM) mode. The column XSELECT HSS XP (150 mm × 2.1 mm × 2.5 μm) (Waters, Milford, MA, USA) with ion-pairing technique was used for the chromatography separation as previously described (38). Peak integration and metabolite quantification were performed using an isotope-ratio-based approach on Multi-QuantTM 3.0.2 (AB SCIEX) software as previously described (39, 40).

## Results

### *P. taiwanensis* VLB120 can easily compensate for single NADH dehydrogenase gene deletions

The NADH dehydrogenase type I operon encoded by *nuoA-N* (PVLB_15600-15660) and the two type II NADH dehydrogenases encoded by *ndh1* (PVLB_13270) and *ndh2* (PVLB_21880) were successfully deleted from the *P. taiwanensis* VLB120 genome using the I-SceI-based pEMG plasmid (28). The double knockout mutants ΔΔ*ndh* and Δ*nuo*Δ*ndh1* were successfully obtained, however several attempts failed to generate the double knockout of *Δnuo* and *Δndh2* (Table 2). All gene deletions were confirmed by Sanger sequencing. The five NADH-dehydrogenase mutants demonstrated that the NADH dehydrogenases, Nuo, Ndh1, and Ndh2 are not essential individually. While the presence of either Nuo or Ndh2 is sufficient to sustain viability of *P. taiwanensis* VLB120, Ndh1 seems to be unable to compensate for the loss of Nuo and Ndh2. Similarly, for *P. aeruginosa* PAO1 it has been reported that single deletions of NADH dehydrogenases result in no growth defect and no decrease in NADH oxidation activity, whereas the double (Δ*nuoIJ*Δ*ndh*) and triple knockout (Δ*nuoIJ*Δ*ndh*Δ*nqrA-F*) had no NADH oxidation activity (41). Concludingly, Nuo and Ndh account for the total NADH dehydrogenase activity in this *Pseudomonas* strain. A likewise total loss of NADH dehydrogenase activity in the obligate aerobic *P. taiwanensis* VLB120 strain seems to be lethal indicating that the strain relies on the presence of NADH dehydrogenases for NADH oxidation and that alternative, native NADH consuming reactions do not suffice to efficiently re-oxidize this vital cofactor under the tested conditions.

**Table 2.** Overview of the NADH dehydrogenase mutants generated in this study. ‘X ‘indicates successful gene deletion.

| Strain | NADH dehydrogenase<br>type I | NADH dehydrogenase type II |  |
| --- | --- | --- | --- |
|  | *nuoA-N* (PVLB_15600-15660) | *ndh1*<br>(PVLB_13270) | *ndh2*<br>(PVLB_21880) |
| Wild type | present | present | present |
| $\Delta ndh1^a$ | present | X | present |
| $\Delta ndh2^a$ | present | present | X |
| $\Delta\Delta ndh^a$ | present | X | X |
| $\Delta nuo^a$ | X | present | present |
| $\Delta nuo\Delta ndh1^a$ | X | X | present |

<sup>a</sup> The megaplasmid pSTY (42) was absent in all mutants.

### The ΔΔ*ndh* mutant showed a growth-phase dependent growth defect

*P. taiwanensis* VLB120 and the five NADH dehydrogenase deletion strains Δ*ndh1*, Δ*ndh2*, ΔΔ*ndh*, Δ*nuo*, and Δ*nuo*Δ*ndh1* were characterized for growth, glucose utilization, CO_2_ formation, and oxygen consumption in batch shake-flask experiments. The single NADH-dehydrogenase type II mutants, Δ*ndh1* and Δ*ndh2*, showed the same growth and sugar co-utilization profile as the wild type *P. taiwanensis* VLB120 (Figure 1A, B, and C). The loss of the megaplasmid pSTY during NADH dehydrogenase deletions resulted in a growth advantage for the generated mutants, which was determined to result in a 14% higher growth rate for *P. taiwanensis* VLB120 pSTY^−^ compared to the pSTY^+^ wild type (data not shown). For a comparison of mutants and wild type, the growth rate of the wild type was corrected accordingly and is referred to as μ_recalc_.

**Figure 1.**
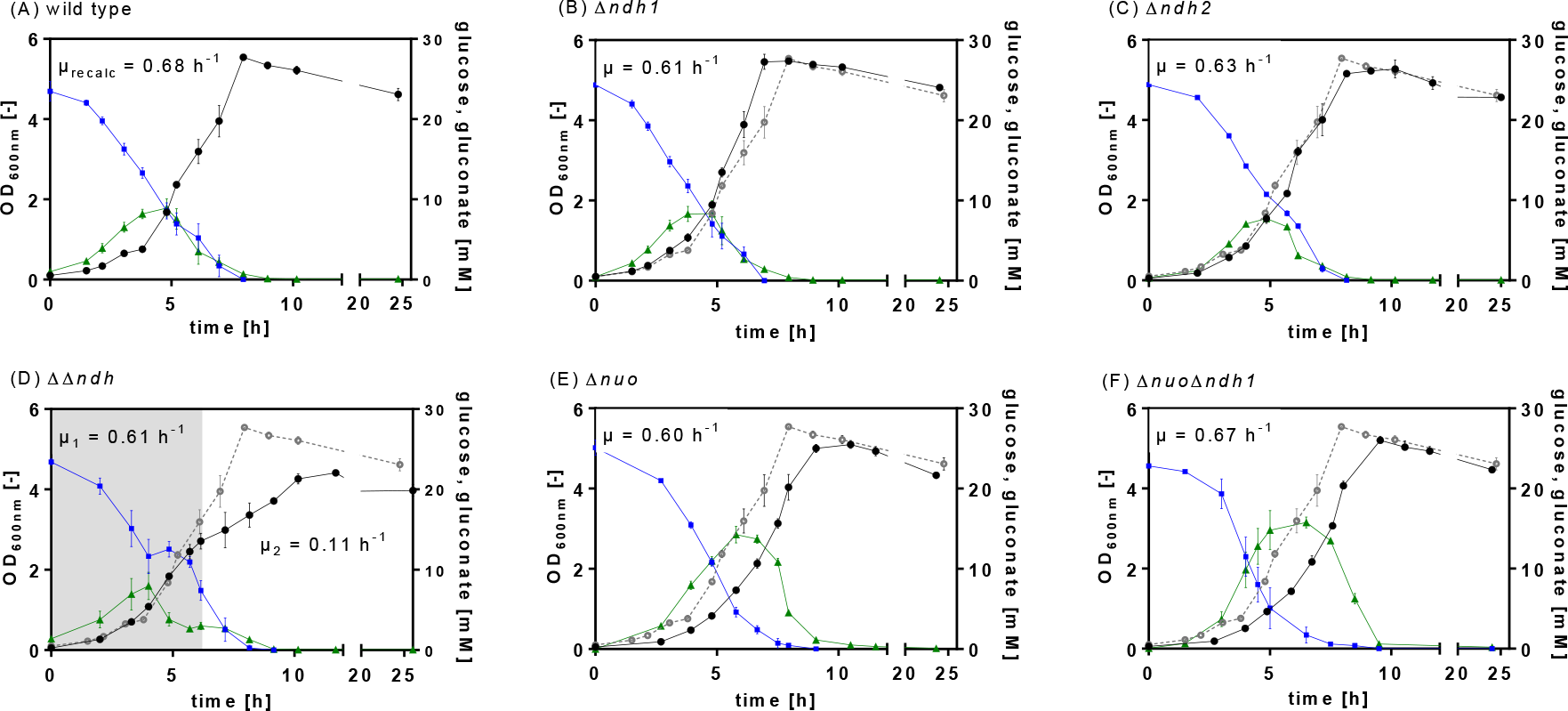
Physiological characterization of *P. taiwanensis* VLB120 (A) and the NADH dehydrogenase deficient mutants Δ*ndh1* (B), Δ*ndh2* (C), ΔΔ*ndh* (D), Δ*nuo* (E), and Δ*nuo*Δ*ndh1* (F). The strains were cultured in MSM with 25 mM glucose. The OD_600nm_ (black circles), glucose (blue squares), gluconate (green triangles) were measured over time. The shadowed area in (D) indicates the first growth phase. The data shown are the mean of biological triplicates; error bars show the standard deviation. μ_recalc_ is the growth rate of *P. taiwanensis* VLB120 pSTY^−^. The wild type OD_600nm_ are plotted (grey, open circles) in graphs (B)-(F) for comparison.

Both NADH dehydrogenases type I mutants Δ*nuo* and Δ*nuo*Δ*ndh1* exhibited wild type-like growth and carbon uptake rates (Figure 1, Table 3). However, the glucose utilization in both type 1 deletion strains differed remarkably. *Pseudomonas* can catabolize glucose either via the phosphorylative or the oxidative pathway. In the latter, glucose is oxidized to gluconate in the periplasm by a membrane-bound glucose dehydrogenase (*gcd*) coupled to the reduction of pyrroloquinoline quinone (PQQ) (Figure 6). The phosphorylative pathway starts in the cytoplasm with the phosphorylation of glucose to glucose-6-phosphate catalysed by the glucokinase (Glk) (43, 44). The type I dehydrogenase mutants, Δ*nuo* and Δ*nuo*Δ*ndh1*, accumulated gluconate to up to 16 mM, twice as high as the concentration of the wild type and all type II deletion strains, Δ*ndh1,* Δ*ndh2*, and ΔΔ*ndh* (Figure 1E-F).

**Table 3.** Calculated carbon uptake rates and gluconate and oxygen consumption of wild type and NADH dehydrogenase mutants during early-exponential growth.

| Strain | Specific carbon uptake rate<sup>a</sup> | Gluconate accumulation | Surplus O<sub>2</sub> consumption |
| --- | --- | --- | --- |
|  | [mmol g<sub>cdw</sub><sup>-1</sup> h<sup>-1</sup>] | [mM]<sup>b</sup> | [mM]<sup>c</sup> |
| wild type | 7.7 | 8.9 ± 0.9 | 9.2 ± 0.9 |
| Δ*ndh1* | 6.9 | 8.6 ± 0.9 | 8.1 ± 0.1 |
| Δ*ndh2* | 7.3 | 7.7 ± 0.2 | 6.3 ± 0.4 |
| ΔΔ*ndh* | 8.8/3.3 | 8.8 ± 0.4 | 7.2 ± 1.2 |
| Δ*nuo* | 7.1 | 16.4 ± 0.4 | 12.9 ± 0.1 |
| Δ*nuo*Δ*ndh1* | 6.6 | 16.1 ± 1.0 | 14.1 ± 2.9 |

<sup>a</sup>For ΔΔ*ndh*, separate growth rates were determined for phase 1 (2-6 h) and phase 2 (6-8 h). For all other mutants, growth rates were calculated for the exponential phase (~3-4 h until ~6-7 h after inoculation); <sup>b</sup>gluconate accumulation was calculated from offline analytic of three biologically independent experiments; <sup>c</sup>the surplus oxygen consumption was calculated from the sectional integrals between the OTR (mmol L<sup>-1</sup> h<sup>-1</sup>) and CTR (mmol L<sup>-1</sup> h<sup>-1</sup>) of the first 6 hours of cultivation (Figure 2); shown is the mean of duplicate experiments.

In contrast, the type II double mutant ΔΔ*ndh* reproducibly showed two growth phases (Figure 1D). After comparable growth to the wild type (Figure 1A), in the mid-exponential growth phase, the growth rate dropped drastically marking the second growth phase. Interestingly, the strong decrease of the growth rate (~86%) was not correlated with an equal reduction of the carbon uptake, which showed a decrease of only ~38%.

Besides the characterization for growth and glucose consumption, the respiratory behavior of the wild type and NADH dehydrogenase mutants was studied (Figure 2, Supplementary Figure S1). Again, only the ΔΔ*ndh* mutant showed a different phenotype characterized by a stagnating oxygen transfer rate (OTR) after 6 hours (Figure 2B). This change in the OTR development is an indication for product inhibition, here, potentially by NADH, which cannot be oxidized at the rate required for fast growth. The onset of the reduced specific oxygen uptake rate also correlated well with the change of the growth rate (Figure 1D). In case of growth on glucose, the expected respiratory quotient (RQ), defined as the ratio of OTR and CTR is close to 1. In contrast to this assumption, the measured OTR for all tested mutants during the first 6 hours of cultivation was higher than the CTR resulting in an RQ below 1 (Figure 2A, Supplementary Figure S1) due to the partial oxidation of glucose to gluconate in the periplasm (Figure 6). Indeed, the surplus of consumed oxygen calculated from the sectional integrals between the OTR and CTR (∫ *OTR dt* - ∫ *CTR dt*), correlated with the produced gluconate (Table 3, Figure 2A). During glucose conversion, roughly half of the overall consumed oxygen was used for the oxidation of glucose to gluconate and the re-oxidation of the reduced PQQ formed by the glucose dehydrogenase activity. Consequently, in the glucose phase, the cells can partially uncouple glucose oxidation and energy provision from NADH formation, relieving the dependence on NADH dehydrogenase activity. The O_2_ and CO_2_ transfer rate (CTR) of Δ*ndh2* and ΔΔ*ndh* (Supplementary Figure S1) showed a double peak, which occurred in the same time frame as glucose depletion, and, hence, might be due to the diauxic shift from glucose to gluconate. We assume that the diauxic shift also occurred in the other strains but was not recorded by the measurement frequency of three measurements per hour.

**Figure 2.**
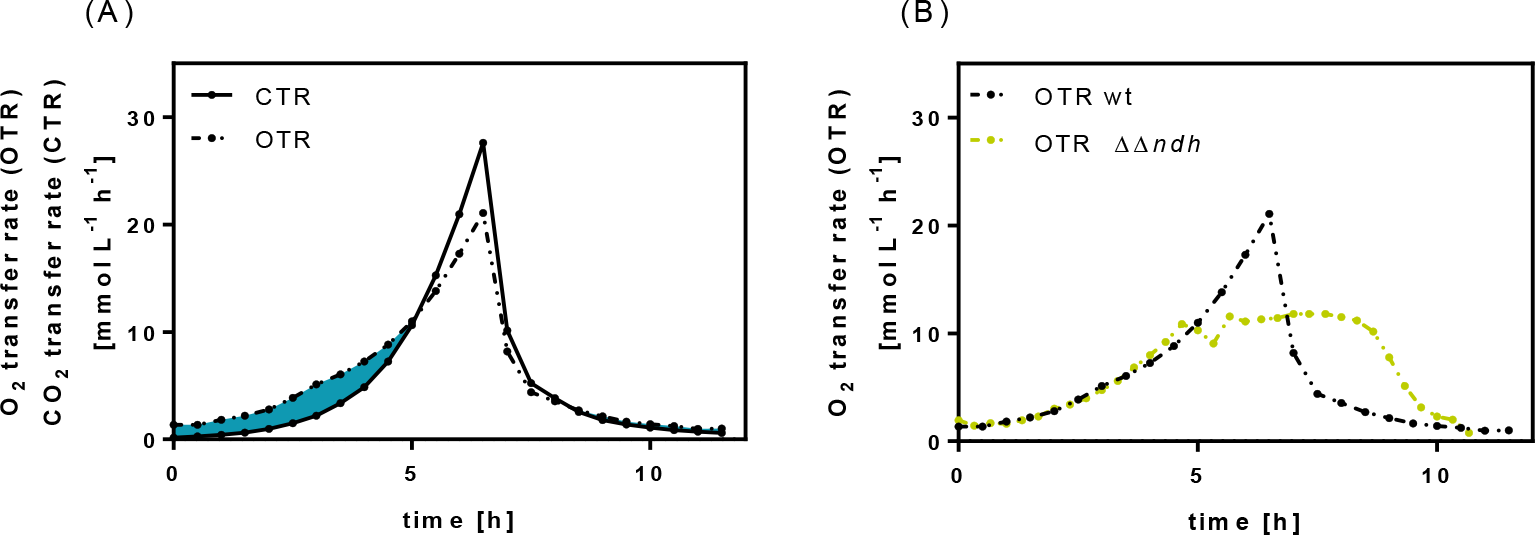
Respiratory activity during a cultivation of *P. taiwanensis* VLB120 and ΔΔ*ndh* mutant. (A) Highlighted area corresponding to the surplus of consumed oxygen, calculated from the sectional integrals between the OTR (dashed line, in mmol L^−1^ h^−1^) and CTR (in mmol L^−1^ h^−1^) on the example of *P. taiwanensis* VLB120.B) Oxygen transfer rates (OTR, dashed line; in mmol L^−1^ h^−1^) during a cultivation of *P. taiwanensis* VLB120 and mutant ΔΔ*ndh.*

### NADH dehydrogenase gene deletions affect expression levels but do not result in altered *in vitro* NADH oxidation activities

To further elucidate the NADH oxidation activity in the different mutants, and hence, the importance of the three NADH dehydrogenases for oxidizing NADH and fueling the electron transport chain, we performed *in vitro* NADH oxidation assays. Inverted membrane vesicles were prepared at early-, mid-, and late-exponential growth phase and the NADH oxidation rate was determined from the decrease of the absorbance at 340 nm over time. As the SDS PAGE showed up to 21 prominent protein bands in the membrane fraction (data not shown), we cannot exclude activity of other membrane-bound NADH-dependent enzymes in the analyzed cytoplasmic fraction, e.g., the transhydrogenase PntAB, which might have contributed to NADH consumption. However, there is a high probability that NADH oxidation is very specific for NADH dehydrogenases as other NADH-dependent enzymes require electron acceptors other than O_2_. In the early-exponential growth phase, in which none of the strains showed a growth defect, all single mutants possessed NADH oxidation activities at levels similar to the wild type of around 1.2 U mg protein^−1^ (Table 4), which is in the range of *in vitro* rates reported for other organisms (45). Overall, the NADH oxidation rate was rather stable in all mutants, indicating a high metabolic flexibility of *P. taiwanensis* VLB120 to maintain redox homeostasis.

**Table 4.**
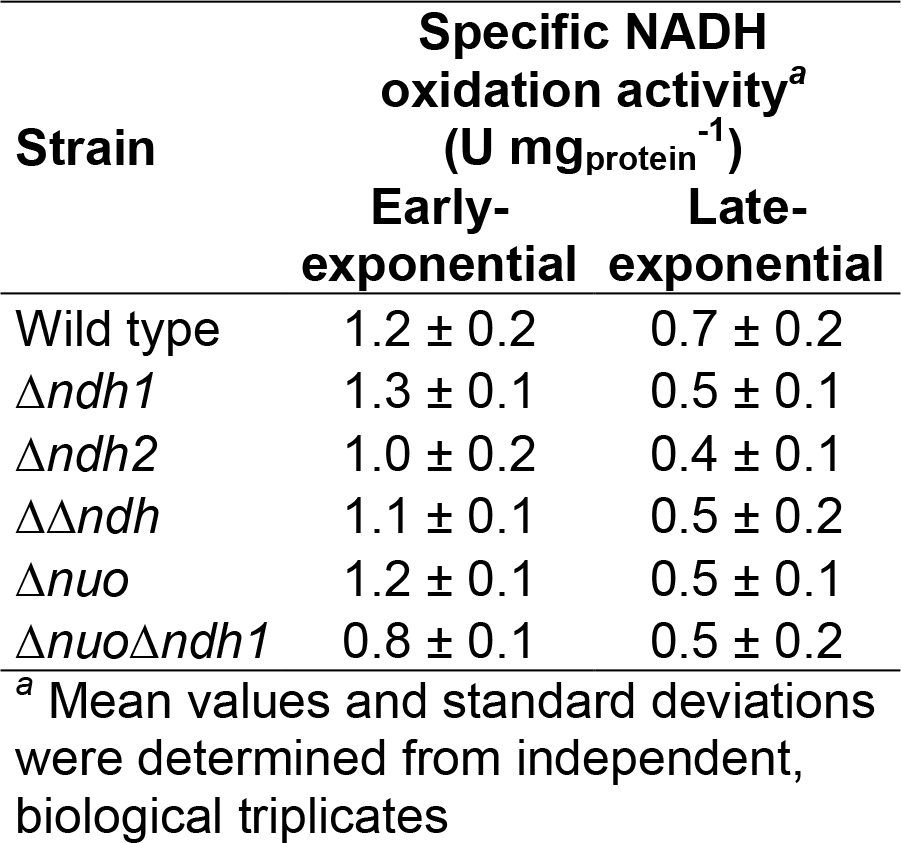
Specific NADH oxidation activities of inverted membrane vesicles of *P. taiwanensis* VLB120 wild type and NADH dehydrogenase mutants at early and late-exponential growth phase.

| Strain | Specific NADH<br>oxidation activity<sup>a</sup><br>(U mg<sub>protein</sub><sup>-1</sup>) |  |
| --- | --- | --- |
|  | Early-exponential | Late-exponential |
| Wild type | 1.2 ± 0.2 | 0.7 ± 0.2 |
| $\Delta ndh1$ | 1.3 ± 0.1 | 0.5 ± 0.1 |
| $\Delta ndh2$ | 1.0 ± 0.2 | 0.4 ± 0.1 |
| $\Delta\Delta ndh$ | 1.1 ± 0.1 | 0.5 ± 0.2 |
| $\Delta nuo$ | 1.2 ± 0.1 | 0.5 ± 0.1 |
| $\Delta nuo\Delta ndh1$ | 0.8 ± 0.1 | 0.5 ± 0.2 |

<sup>a</sup> Mean values and standard deviations were determined from independent, biological triplicates

**Table 5.**
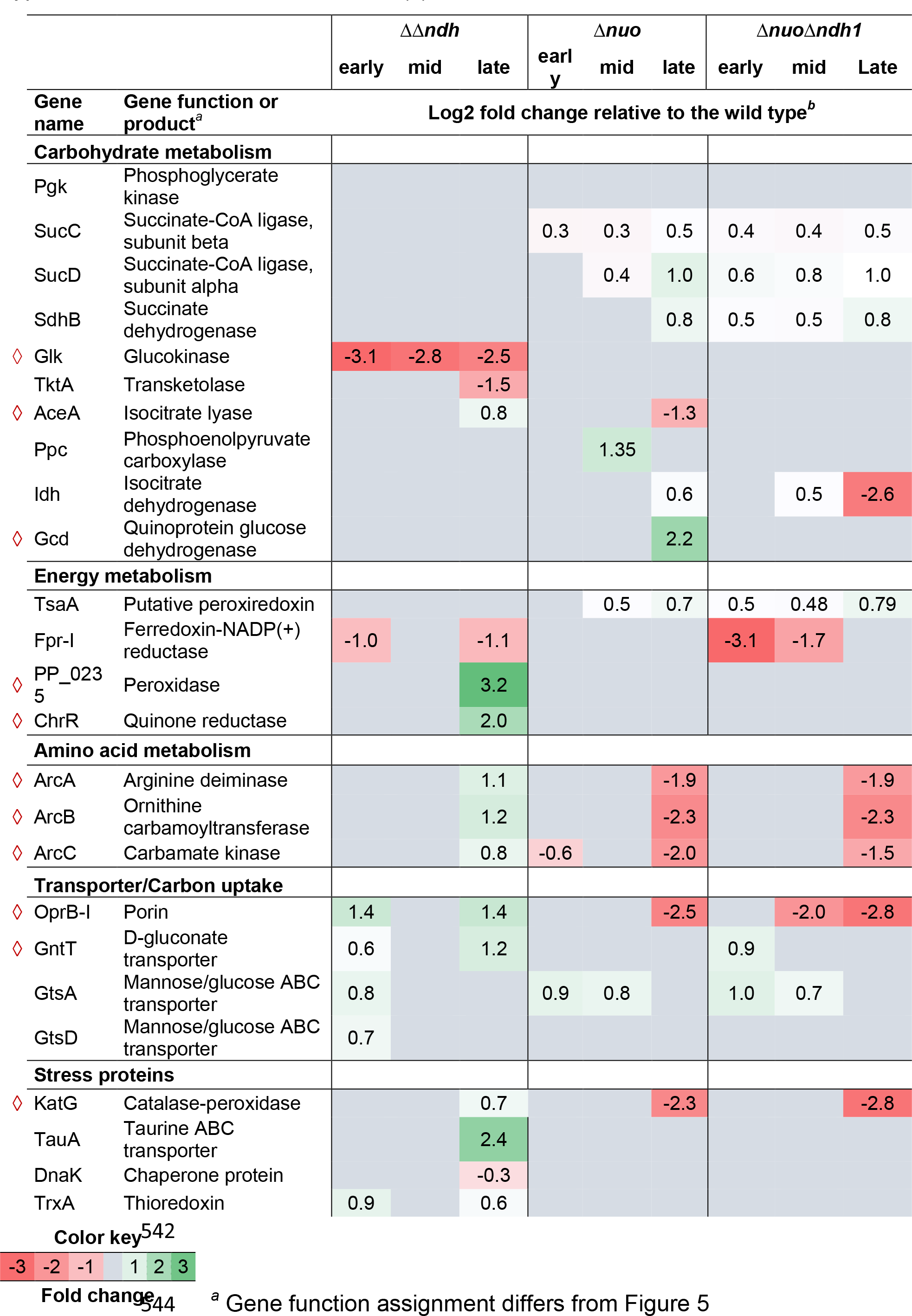
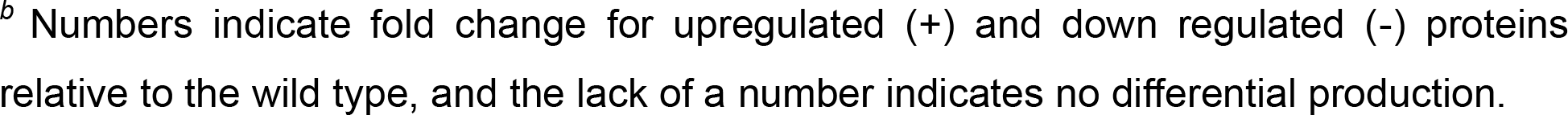
Protein abundance in NADH dehydrogenase mutants in comparison to the wild type. Proteins marked with a diamond (◊) are discussed in the text.

To further substantiate this hypothesis, we examined potential changes at the transcriptional level by qPCR on samples taken in the early, mid-, and late-exponential growth phase. HPLC analysis showed that glucose and/or gluconate were still left when sampling the late-exponential growth phase, i.e., the cells were still metabolically active (data not shown). The fold changes were normalized against the wild type in the corresponding growth phase. The single and double deletions of NADH dehydrogenases type II encoding genes (Figure 3A-C) had only minor effects (max. fold change of 1) on the remaining NADH dehydrogenase gene expression. While the type I deletions strains, Δ*nuo* and Δ*nuo*Δ*ndh1*, showed a substantial upregulation of the *ndh2* gene expression (Figure 3D-E), the expression of the *ndh1* gene in both the Δ*nuo* and Δ*nuo*Δ*ndh1* was unaffected; we only observed a small increase for mutant Δ*nuo* in the early growth phase. This finding suggests that *ndh2* is probably the only NADH dehydrogenase gene that is regulated by the cellular NADH economy. The consequent essentiality would further explain why the double deletion of *nuo* and *ndh2* was lethal. The observation shows that ΔΔ*ndh* is only growth impaired in the mid-to late-exponential phase. This indicates that either the Nuo complex is less active in the late growth phase or that the PQQ-dependent glucose dehydrogenase activity during the early growth phase enables sufficient ATP synthesis independent of NADH dehydrogenase activity.

**Figure 3.**
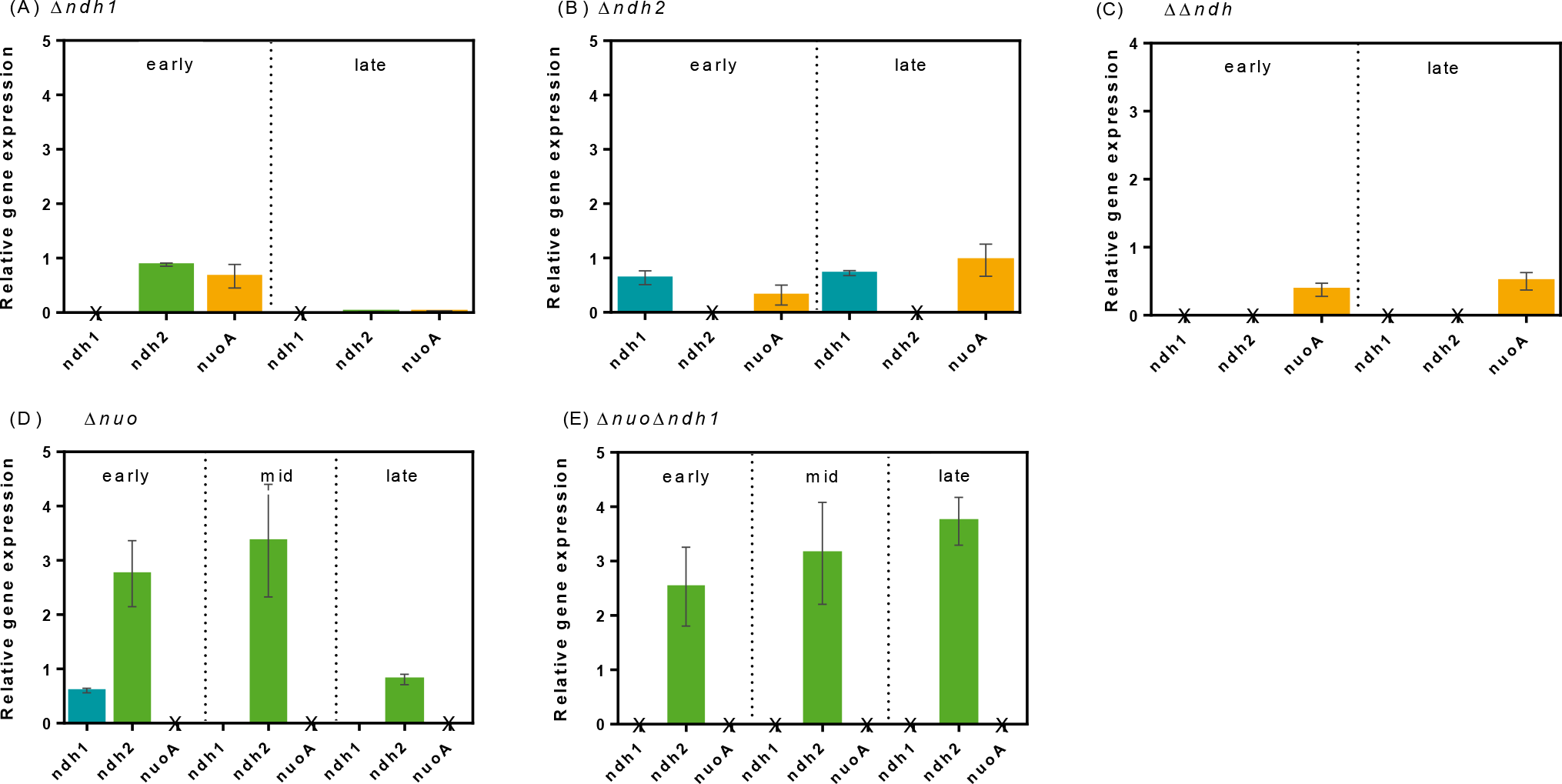
Relative gene expression of the NADH dehydrogenase encoding genes *ndh1*, *ndh2*, and *nuoA* in NADH dehydrogenase mutants Δ*ndh1* (A), Δ*ndh2* (B), ΔΔ*ndh* (C), Δ*nuo* (D), and Δ*nuo*Δ*ndh1* (E) at early-, mid-, and late-exponential growth phase normalized to the corresponding values of the wild type. mRNA abundance was determined by quantitative PCR. Values were normalized to the relative transcript level of *P. taiwanensis* VLB120 in the corresponding growth phase. *nuoA* was used as a proxy for the expression of the *nuo* operon. Gene deletions in the respective mutant are marked with ‘X’ and were not analyzed by qPCR. Experiments were performed in biological triplicates.

### Double deletion of the type II NADH dehydrogenases affect intracellular redox cofactor levels

We found that the NADH oxidation rate was not or only slightly compromised by the introduced gene deletions but that the *ndh2* level was significantly upregulated suggesting that its expression is controlled by the NADH level. For that reason, we determined the intracellular abundance of NADH and NAD^+^ in the early-, mid-, and late-exponential growth phase. Since the two single mutants of type II had no growth phenotype and showed no obvious changes on the transcriptional level, we restricted the analysis to the two double mutants and single *nuo* deletion mutant Δ*nuo*, ΔΔ*ndh,* and Δ*nuo*Δ*ndh1*.

The Δ*nuo*Δ*ndh1* showed a higher NADH/NAD^+^ ratio in the late-exponential growth phase but also high variability in the triplicate experiments curtailing the statistical significance. The double mutant ΔΔ*ndh* had a significantly increased NADH/NAD^+^ ratio in the mid- and late-exponential growth phase compared to the wild type (Figure 4). This significantly increased NADH/NAD^+^ ratio in ΔΔ*ndh* might have triggered the observed growth change in the mid-exponential growth phase.

**Figure 4.**
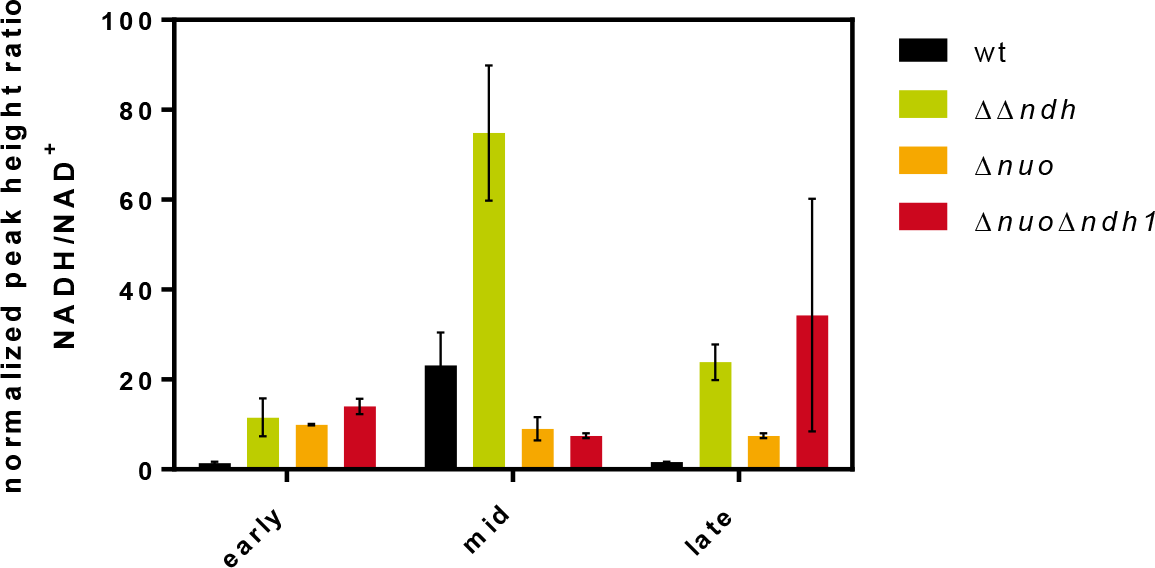
Quantification of the NADH/NAD^+^ ratio in the *P. taiwanensis* VLB120 and the NADH dehydrogenase mutants ΔΔ*ndh*, Δ*nuo*, and Δ*nuo*Δ*ndh1* in early-, mid- and late-exponential growth phase.

### Proteomic analysis reveals rerouting of the carbon flux in the ΔΔ*ndh* mutant

We further performed shotgun proteomic analysis to explain possible metabolic changes in early-, mid-, and late-exponential growth phase in *P. taiwanensis* VLB120 due to NADH dehydrogenase deletions. The relative quantitative results were used to categorize the detected proteins into three groups: (1) Significantly upregulated or (2) downregulated proteins (fold change > 2, adjusted p-value < 0.05), and (3) weak/no effect proteins (fold change < 2). The proteins were further grouped into functional categories according to the KEGG database classification (46), e.g., transport, carbohydrate metabolism, amino acid metabolism, Supplementary Table S 2) the most strongly represented categories are summarized in Figure 5.

**Figure 5.**
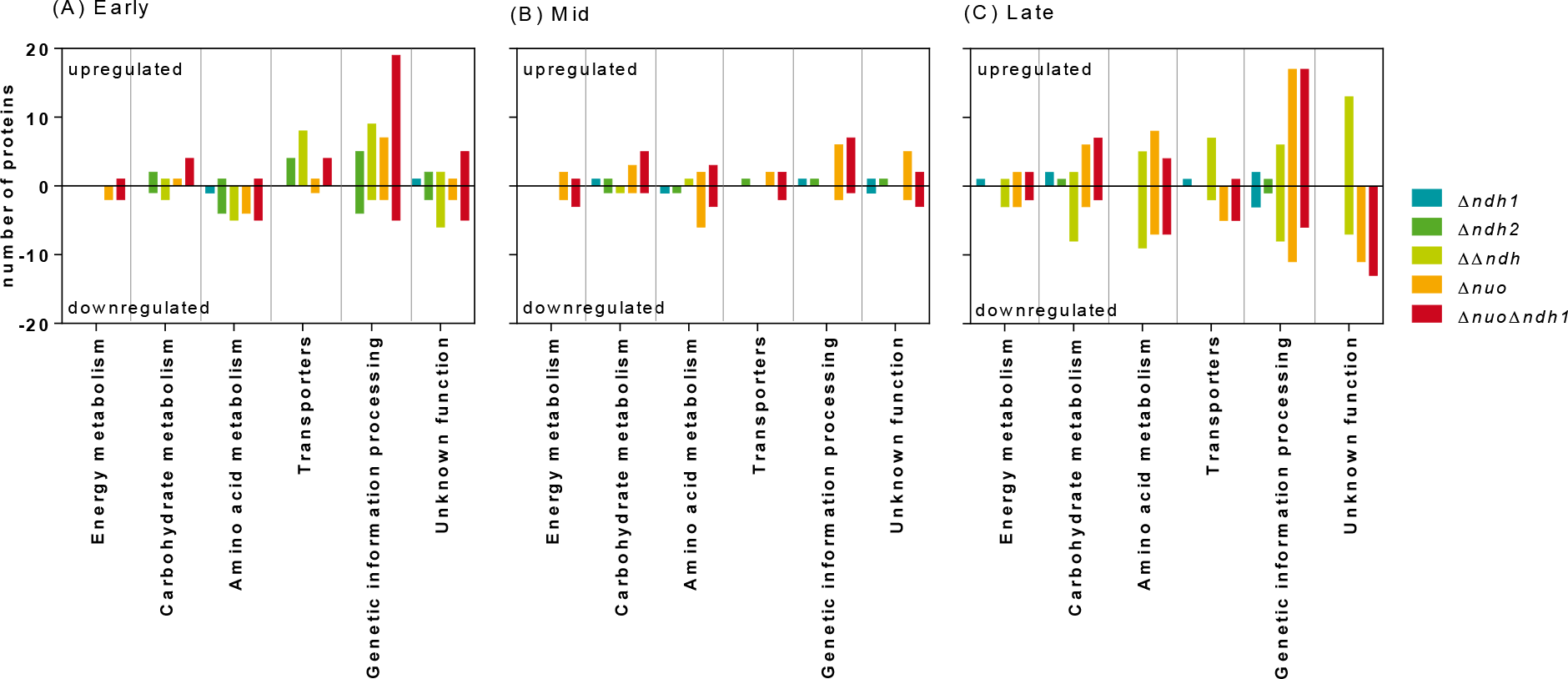
Significant changes at proteome level of *P. taiwanensis* VLB120 NADH dehydrogenase mutants in early- (A), mid- (B), and late-exponential (C) growth phase relative to the wild type. Proteins are clustered into functional categories according to the KEGG classification system (46). Each bar represents the number of proteins in the depicted category the abundance of which was either increased or decreased in response to NADH dehydrogenase deficiency. Experiments were performed in biological triplicates.

In accordance with the physiological and transcript data, we did not observe major changes in the proteome for either NADH dehydrogenase type II single mutants (Δ*ndh1*: 9 of 24 proteins significantly up/downregulated, Δ*ndh2*: 8 of 36 proteins significantly up-/downregulated; Figure 5, Supplementary Table S2, Supplementary File S3). Proteomic changes in both type I mutants (Δ*nuo*: 50 of 139 proteins significantly up-/downregulated, Δ*nuo*Δ*ndh1*: 60 of 165 proteins significantly up-/downregulated) were more significant compared to the type II single gene knockout mutants and very similar to each other (Figure 5). The double deletion mutant ΔΔ*ndh* showed more alterations in the proteome in the early- and late-exponential phase (17 and 37 of 107 proteins significantly up-/downregulated) than in the mid-exponential phase (2 significantly up-/downregulated proteins) (Figure 5 and Supplementary Table S2).

In the following paragraphs we are focusing on changes observed the ΔΔ*ndh* mutant for proteins related to carbon uptake, energy generation, and oxidative stress response and highlight particular differences to the type 1 NADH dehydrogenase mutants.

The OprB-I porin (PVLB_20075), a carbohydrate selective porin, and the D-gluconate transporter GntT (PVLB_13665) located in the outer and inner membrane, respectively, were higher increased in the ΔΔ*ndh* mutant during the early- and late-exponential growth phase, while the glucokinase quantity was strongly reduced in all growth phases. These data suggest that ΔΔ*ndh* oxidized glucose via glucose dehydrogenase (Gcd) to gluconate to a greater extent than the wild type. In contrast, the quantity of OprB-I in the type I NADH dehydrogenase mutants during the later growth phase was decreased. This might, however, be explained by the faster glucose depletion in these mutants (Figure 1).

During the late-exponential growth of the ΔΔ*ndh* mutant, all enzymes of the arginine deiminase (ADI) pathway were more strongly expressed while they were significantly downregulated in the NADH dehydrogenase type I mutants (Table 6). This pathway catalyzes a three-step conversion of arginine to ornithine, ammonium, and carbon dioxide coupled to ATP generation (47). Likewise, the isocitrate lyase (AceA), the first enzyme of the glyoxylate shunt, was upregulated in the ΔΔ*ndh* mutant but downregulated in the Δ*nuo* mutant, which rather showed a slight upregulation of the 2-oxoglutarate dehydrogenase complex of the TCA cycle during mid- and late-exponential growth. These changes indicate that the mutant ΔΔ*ndh* used the glyoxylate shunt and not exclusively the TCA cycle in the late-exponential growth phase.

We further observed remarkable changes of proteins combating oxidative stress. While deletion of the *nuo* operon (Δ*nuo* and Δ*nuo*Δ*ndh1)* resulted in a generally reduced abundance of proteins involved in the oxidative stress response, those mutants deficient in one of the two type II dehydrogenases displayed increased levels of peroxidases and peroxiredoxin proteins. The abundance of the catalase-peroxidase KatG was extremely decreased in both type I mutants whereas it was weakly increased in the ΔΔ*ndh* mutant. Additionally, the peroxidase encoded by PP_0235 and the quinone reductase ChrR were only higher expressed in the ΔΔ*ndh* mutant with the latter being reported to be induced by superoxide (48) while peroxiredoxin AhpC was weakly upregulated in the single gene deletion mutants, Δ*ndh1* and Δ*ndh2* (Supplementary File S3). These findings indicate that the deletion of the *nuo* complex reduces oxidative stress while it is increased in the mutants deficient in type I dehydrogenase.

## Discussion

The presented in-depth analysis of NADH dehydrogenase mutants revealed a high metabolic robustness of *P. taiwanensis* VLB120 to partial loss of the three NADH dehydrogenases, but also the essentiality of residual NADH dehydrogenase activity, as the simultaneous deficiency of Nuo and Ndh2 was lethal likely due to inefficient NADH oxidation or ATP provision.

In accordance with the observed phenotypic robustness, NADH oxidation activities in the mutant strains were not reduced. While this can be explained for those mutants deficient in the *nuo* operon with a significant upregulation of *ndh2*, no transcriptional changes of NADH dehydrogenase related genes were observed for the other mutants. In contrast the mutant with Nuo as the sole NADH dehydrogenase (ΔΔ*ndh*) showed a growth phenotype upon mid-exponential growth phase, accompanied by various metabolic changes summarized in Figure 6. The wild type like growth of the ΔΔ*ndh* mutant on glucose is likely sustained by switching from glucose phosphorylation to direct oxidation in order to partially uncouple the oxidation of the carbon source from NADH formation.

**Figure 6.**
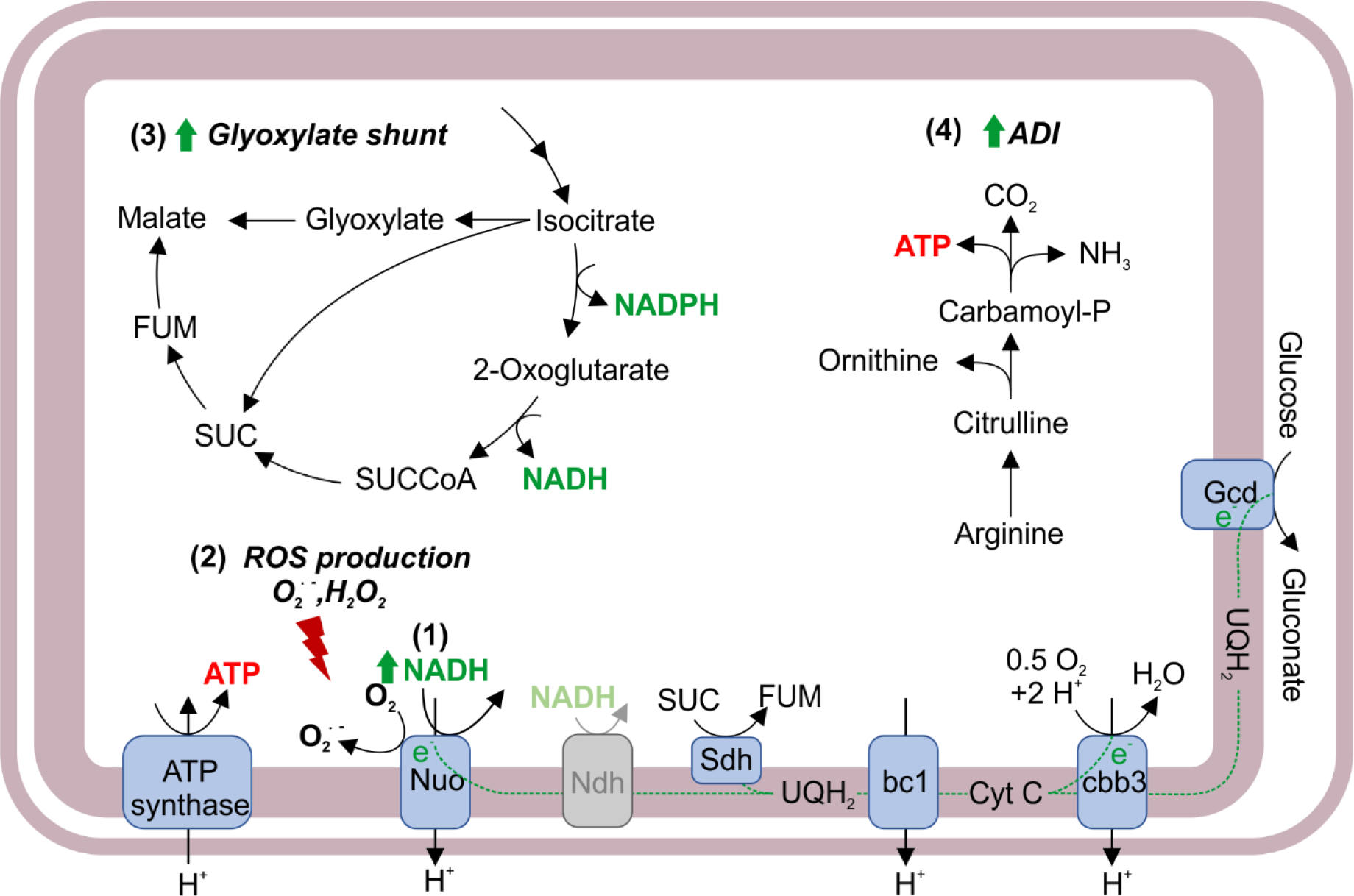
Putative rerouting of the carbon flux caused by type II NADH dehydrogenase deficiency in *P. taiwanensis* VLB120. An increased NADH/NAD^+^ ratio ****(1)**** triggers ROS production ****(2)**** by NADH dehydrogenase Nuo activity (52, 53). The corresponding cell response to oxidative stress is rerouting the carbon flux through the TCA cycle into the glyoxylate shunt ****(3)**** to reduce NADH formation (56–58) and scavenging of reactive oxygen species by glyoxylate (56, 59). Limited ATP provision from oxidative phosphorylation might be mitigated by upregulation of the ADI pathway (*4*)(65, 66). The light representation of the Ndh dehydrogenase indicates deficiency of both isozymes. ETC, electron transport chain; ROS, reactive oxygen species; ADI, arginine deiminase pathway; Nuo, type I NADH dehydrogenase; Ndh, type II NADH dehydrogenase; Sdh, succinate dehydrogenase; bc1, cytochrome bc1 (complex III); cbb3, cytochrome cbb3 (complex IV); QH_2_, ubiquinol; Q, ubiquinone; SUC, succinate, SUCCoA, succinyl-CoA; FUM, fumarate.

*In vitro* studies have shown the formation of reactive oxygen species (ROS) such as superoxide (O_2_^·−^) and hydrogen peroxide (H_2_O_2_) by enzymes of the electron transport chain due to electron leakage to oxygen. Nuo (complex I) and cytochrome bc1 (complex III) are considered the main sites for ROS in mitochondria and *P. fluorescens* (49, 50) but rather the type 2 dehydrogenase in, e.g., *E. coli* 51. It has further been reported that an oversupply of NADH can enhance ROS production (52, 53), which activates the cellular antioxidant defense system and can lead to cell death (49, 54). Due to a relatively high NADH/NAD^+^ ratio in the mid-exponential growth phase, ΔΔ*ndh* might have been faced with higher ROS production than the wild type. This hypothesis is supported by the RAMOS data (Figure 2), which indicated product inhibition after six hours potentially caused by high NADH levels or oxidative stress. In line with this hypothesis, it has been shown that *M. tuberculosis* NADH dehydrogenase mutants with a similarly elevated NADH/NAD^+^ ratio were more susceptible to (additional) oxidative stress than those with a lower NADH/NAD^+^ ratio (55). Increased ROS production due to an increased NADH/NAD^+^ ratio could be one reason for the strongly reduced growth rate in the later growth stages and the trigger for metabolic rerouting in ΔΔ*ndh* (Figure 6).

The activation of the glyoxylate shunt as indicated by the proteome data might further contribute to stress reduction in two ways. Firstly, this shortcut of the TCA cycle bypasses NADH-producing steps, effectively attenuating NADH generation (56–58). Secondly, the glyoxylate formed by the isocitrate lyase AceA activity of this pathway, upregulated in ΔΔ*ndh*, might react with hydrogen peroxide to produce formate and CO_2_ (56, 59). This ROS combating strategy was also reported in *Pseudomonas aeruginosa*, *Burkholderia cenocepacia*, and *Staphylococcus aureus*, even though *S. aureus* has no functional glyoxylate shunt (56, 60–62). Note, however, that neither higher formate dehydrogenase abundance nor formate accumulation was observed.

ROS production seems to be not elevated in the *nuo* mutants with supposedly higher Ndh2 activity because our data showed no activation of any of these mechanisms. The same was suggested for corresponding *M. tuberculosis* mutants (55).

A probable energy shortage due to reduced respiratory activity might have been counteracted in ΔΔ*ndh* by activation of the ADI pathway as an alternative ATP source. This pathway generates 1 mol ATP per mol arginine (47, 63) and was described in *P. aeruginosa* to provide ATP under oxygen limiting conditions (47). We exclude oxygen limitation as an inducer of the ADI pathway in *P. taiwanensis* VLB120 under the conditions tested here since the pathway was only expressed in the ΔΔ*ndh* mutant, although all analyzed strains were grown under the same conditions and had similar oxygen uptake rates. Rather, energy depletion might have activated the ADI pathway in ΔΔ*ndh* on gluconate as described in lactic bacteria (64) or other *Pseudomonas* strains (65), e.g., in *P. putida* DOT-T1E under energy-demanding solvent stress conditions (66, 67).

In this study, we showed a high metabolic flexibility of *P. taiwanensis* VLB120 to interventions in the redox metabolism, which confers robust phenotypic behavior by rerouting of metabolic fluxes. This metabolic adaptability and phenotypic robustness can be advantageous for biocatalysis but simultaneously be challenging because it impedes the prediction of mutant behavior and can lever out metabolic engineering efforts. Hence, to effectively turn this promising microbe into a controllable, biotechnological workhorse, further systems biological and physiological analyses are needed.

## Acknowledgement

This study has been conducted within the ERA SynBio project SynPath (Grant ID 031A459) with financial support of the German Federal Ministry of Education and Research and is part of the Joint BioEnergy Institute, supported by the Office of Science, Office of Basic Energy Sciences and Office of Biological and Environmental Research of the U.S. Department of Energy under Contract No. DE-AC02-05CH11231. BEE and SCN acknowledge financial support by the German Academic Exchange Service (DAAD) through the thematic network Aachen-California Network of Academic Exchange (ACalNet) funded by the German Federal Ministry of Education and Research (BMBF). LMB acknowledges funding by the Cluster of Excellence “The Fuel Science Center - Adaptive Conversion Systems for Renewable Energy and Carbon Sources” (EXC 2186), which is funded by the Excellence Initiative of the German federal and state governments to promote science and research at German universities. We thank Stephani Baum and Uwe Conrath for sharing lab equipment and for technical support. We are grateful to Itay Budin for discussions and instruction on the membrane isolation and *in vitro* assays and Sophia Nölting for support with qPCR measurements. We thank Benedikt Wynands and Nick Wierckx for sharing the strain *P. taiwanensis* VLB120 pSTY^−^ and Victor de Lorenzo (Centro Nacional de Biotechnología - CNB, Madrid) for providing plasmid pEMG.

